# Lipid-Based Transfection of Zebrafish Embryos: A Robust Protocol for Nucleic Acid Delivery

**DOI:** 10.1101/2024.04.11.589140

**Authors:** Aslihan Terzi, Tiger Lao, Adrian Jacobo

## Abstract

Zebrafish, a widely used model organism in developmental and biomedical research, offers several advantages such as external fertilization, embryonic transparency, and genetic similarity to humans. However, traditional methods for introducing exogenous genetic material into zebrafish embryos, particularly microinjection, pose significant technical challenges and limit throughput. To address this, we developed a novel approach utilizing Lipofectamine LTX for the efficient delivery of nucleic acids into zebrafish embryos by lipid-based transfection. Our protocol bypasses the need for microinjection, offering a cost-effective, high-throughput, and user-friendly alternative. This protocol outlines new strategies for gene delivery in zebrafish to enhance the efficiency and scope of genetic studies in this model system.

## Introduction

Following George Streisinger’s aspirations and work to uncover the genetic logic of neural development in the 1960s, zebrafish have become one of the premiere research organisms used to investigate the genetic and molecular underpinnings of various biological processes in vertebrates. Zebrafish are now a powerful and versatile model in scientific research due to their external fertilization and development, embryonic and larval transparency, genetic similarity to humans, and amenability to genetic manipulations.

Delivering exogenous DNA or RNA into cells is a powerful tool for studying gene function. Gene delivery methods in zebrafish can generate transient products in cells or lead to alterations in genomes that persist through generations. Common applications of gene delivery include the expression of transient payloads, gene knock-down, -out or -in, and transposon or integrase-mediated genome integration of transgenic constructs (Auer et al., 2014, Jao et al., 2013, Kwan et al., 2007, Medishetti et al., 2022, Mi and Andersson, 2023, Lalonde et al., 2023).

Microinjection has been the primary nucleic acid delivery method since the emergence of zebrafish as a model for genetic studies. This method is highly efficient, yet technically demanding, time-consuming, and labor-intensive, creating a major bottleneck that prevents high throughput generation of genetically modified lines for forward genetic screens. Microinjection also requires specialized equipment, including a dissection scope, a micromanipulator, and a picopump (Pei and Burgess, 2019, Rosen et al., 2009). Furthermore, microinjection needles must be trimmed to a sharp point for optimal chorion and embryo penetration, adding an extra variable step to generating good and consistent needles for injections (Pei and Burgess, 2019). Moreover, the genetic material needs to be carefully loaded into the needle to avoid clogging and air bubbles. If injecting multiple unique constructs in one setting, it is necessary to repeat this tedious process every time. MIC-Drop has been developed to inject intermixed droplets with target genes for large-scale genetic screens in a single needle, however, it does not overcome the need for an injection instrument and laborious process (Parvez et al., 2021). Even with the correct instruments, mastering injections necessitates technical expertise only acquired by practice over time. Finally, when the goal is to create genetically modified embryos, these need to be microinjected at the one or two-cell stage to minimize mosaicism, which leaves a narrow window of time to execute this procedure after egg fertilization. Because of these constraints, the number of different genetic constructs that can be delivered in a single injection session is limited to a handful at most.

Electroporation has been tested in zebrafish as an alternative to microinjection. During electroporation, an electric pulse forms pores in the cell membrane, facilitating nucleic acid entry (Kumar et al., 2019). This method is commonly used to control temporal gene delivery after embryonic development in zebrafish, which enables analysis of later developmental stages and adult tissues (Durovic and Ninkovic, 2019, Kera et al., 2010, Thummel et al., 2011). Successful electroporation at earlier stages of embryonic zebrafish has been reported only once and was performed at the eight-cell stage (Zhang et al., 2020). The electroporation protocol requires large amounts of DNA (20 µg) and can only be performed on 15 to 20 embryos at a time. Furthermore, it necessitates special equipment, and detailed optimization to find the best parameters for nucleic acid delivery.

Another alternative for microinjection is lipid-based transfection, or lipofection, which is cost-effective, simple, and high throughput. Lipofection employs a positively charged liposome to encapsulate DNA or RNA and relies on endocytosis for delivery into cells (Felgner et al., 1987, Zhu and Mahato, 2010, Pemberton et al., 2024). This method bypasses the intracellular trafficking and the endosomal pathways, leading to higher transfection efficiency (Cardarelli et al., 2016). Here, we developed a protocol for gene delivery in zebrafish via lipofection with Lipofectamine LTX. This protocol is straightforward, easy to follow, overcomes time constraints, and does not require special equipment or technical expertise. It can be used with multiple constructs in parallel, to accelerate the delivery of nucleic acids in zebrafish. To our knowledge, this is the first time that lipid-based transfection is used with zebrafish embryos, obviating the need for microinjection.

## Methods

### Animal care

Care and experimental procedures for zebrafish adhere to the protocols approved by the institutional animal care and use committee at the University of California San Francisco (UCSF). The fish were bred and maintained at 28°C and were fed thrice daily using an automatic feeder (Lange et al., 2021). Embryos were collected 15 minutes (min) after breeding. Transfected embryos were maintained in E3 medium with methylene blue at 28°C.

### mRNA preparation

The coding sequence (GenBank LC601652.1) for Stay-Gold (Hirano et al., 2022) was codon-optimized for zebrafish using Integrated DNA Technology’s (IDT) online tool and synthesized as a gBlock gene fragment by IDT (Coralville, IO). This fragment was cloned into a pCS2+ backbone and transcribed in vitro using mMessage mMachine SP6 kit (Thermo Fisher Scientific, Waltham, MA). Zebrafish codon optimized Tol2 mRNA was synthesized by RiboPro (Oss, The Netherlands).

### Embryo transfection

A detailed protocol is provided as supplementary material. Briefly, embryos were dechorionated in 0.5 mg/mL pronase (Sigma-Aldrich, St. Louis, MO) until the chorion exhibited wrinkling (Mito et al., 2020). Embryos were then washed with 1X E3 at least four times to remove pronase and chorions. Dechorionated embryos (n=30) were placed in 30mm glass dishes filled with 1x E3 media using flame-polished glass pipettes.

mRNA-only transfection: In a microcentrifuge tube, 0.5 to 2 µg of StayGold mRNA was mixed with Plus™ reagent at a 1:1 ratio (µg:µL) in optiMEM (Thermo Fisher Scientific, Waltham, MA) to a final volume of 50 µL and incubated for 5 min at room temperature.

mRNA and plasmid co-transfection: In a microcentrifuge tube, 2 µg of Tol2 plasmid and 1 µg of Tol2 mRNA were mixed with Plus™ reagent 1:1 ratio (µg:µL) in optiMEM and incubated for 5 min at room temperature. In a different tube, 3 µL of Lipofectamine LTX (Thermo Fisher Scientific, Waltham, MA) was diluted in optiMEM. Plasmid-mRNA-Plus reagent mix (50 µL) was added onto Lipofectamine LTX (50 µL), mixed by finger flicking, and incubated for at least 5 min at room temperature. In a different tube, 1-2 µL of Lipofectamine LTX was diluted in optiMEM to a final volume of 50 µL. mRNA-Plus reagent mix (50 µL) was added onto Lipofectamine LTX (50 µL), mixed by finger flicking, and incubated for at least 5 min at room temperature.

400 µL of 1X E3 (without methylene blue) was added to the Lipofectamine-nucleic acid mix before adding it to the zebrafish embryos. Embryos were then incubated at 28°C for 1-4 hours. After incubation, 1X E3 with methylene blue was added to the media, and embryos were transferred into 60 mm glass dishes with fresh 1X E3/methylene blue for further development.

### RNA isolation and RT-qPCR

Embryos were anesthetized with 0.016% tricaine methanesulfonate (Pentair, Apopka, FL) and collected in 1.5 mL microcentrifuge tubes at 1 day-post-transfection. 10-15 embryos were lysed and homogenized in 0.5 mL of TRI-Reagent (Zymo Research, Irvine, CA) with a pellet pestle. Homogenized embryos were used for total RNA isolation with Direct-zol RNA Microprep (Zymo Research, Irvine, CA). Primers against StayGold mRNA were designed using PrimerBlast (Ye et al., 2012). 50 ng of total RNA was used with Luna Universal One-Step RT-qPCR (NEB, Ipswich, MA) to detect StayGold mRNA expression. Samples were run in BioRad CFX96 Real-Time System (Hercules, CA) for 35 cycles. For each condition, samples were analyzed in three independent wells in a single 96-well plate. The experiment was repeated at least in three biological replicates of total RNA isolated from independent cohorts of embryos. The 2^∆∆*Ct*^ method was used to calculate the relative gene expression in CFX Maestro software, using rpl13a as a reference gene (Weaver et al., 2016), and relative expression was normalized against 1 µg-transfected samples.

Primers used:

> rpl13a forward: 5’-*TCCTCCGCAAGAGAATGAAC*-3’,
>
> rp13a reverse: 5’-*TGTGTGGAAGCATACCTCTTAC*-3’,
>
> StayGold forward: 5’-*GGCGATATGAACGTGTCCCT*-3’,
>
> StayGold reverse: 5’-*GCCTCCACGCACTTACTCTT*-3’.

### Image Acquisition and Analysis

Embryos were anesthetized in 0.016% tricaine methane-sulfonate (Pentair, Apopka, FL) and mounted in 3% methylcellulose (Sigma-Aldrich, St. Louis, MO). Embryos were imaged using Leica Thunder Imager M205 FCA, a 1x Plan APO objective, and a Leica K5 sCMOS camera. Images were analyzed using FIJI (Schindelin et al., 2012). Fluorescence intensity was measured inside ROIs containing whole embryos, manually drawn using the free-hand tool. As background, a rectangular area close to the embryo was set as ROI. The background value was subtracted from the whole-fish ROI. The resulting measurements were normalized using embryos transfected with 1µg of mRNA as the reference value.

### Statistical Analysis

Data sets were analyzed using GraphPad Prism software version 10.2.0 (GraphPad Prism, RRID:SCR-002798; La Jolla, CA). Student’s t-test (two-tailed test) was used to identify differences among means for data sets containing two groups. For data sets containing three or more groups, one-way ANOVA or Kruskal-Wallis tests were used to identify differences among means. If a significant difference was detected with the ANOVA, the Tukey HSD or Sidak correction method was used for multiple comparisons among groups. If a significant difference was detected in the Kruskal-Wallis test, Dunn’s multiple comparisons test was used. p values <0.05 were considered significant. Graphs show the mean *±* standard error of the mean (SEM) for each group. Data were averaged from at least three independent experiments.

## Results

### mRNA can be effectively delivered into zebrafish embryos with lipid-based transfection

To test the protocol, we transfected in vitro transcribed StayGold (Hirano et al., 2022) mRNA into one-cell stage zebrafish embryos. Across multiple experiments we observed that 95% of the embryos showed strong fluorescence (Fig.1A-C), demonstrating that mRNA transfection is highly efficient. Lipofectamine did not show toxicity to embryos at the required concentrations. We did not observe any difference in survival rates of transfected and untransfected controls up to 5dpt (Fig. 1D). Moreover, mRNA-transfected embryos developed normally (Fig. 1D). Overall, we found that Lipofectamine is a safe and effective alternative to microinjection in zebrafish embryos.

**Fig. 1.**
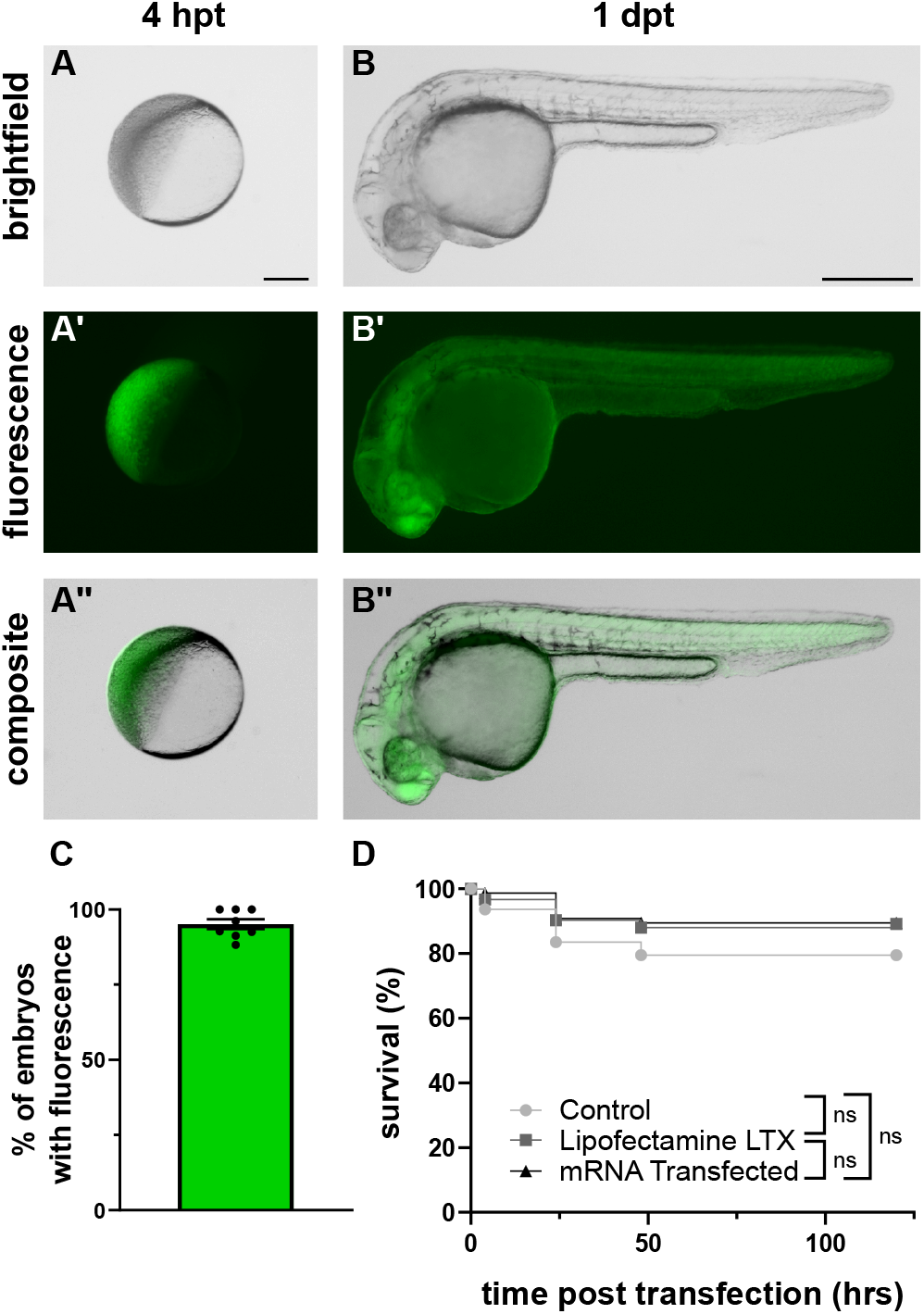
Efficient delivery of mRNA into one-cell-stage zebrafish embryos with lipofection. (A-A”) Embryo transfected with StayGold mRNA at 4 hours post-transfection (hpt) and (B-B”) 1-day post transfection (dpt). (C) The transfection efficiency is shown as the percentage of embryos with fluorescence. The graph shows the mean *±* SEM. (D) Survival of untransfected control, Lipofectamine (LTX) transfected control, and mRNA transfected embryos at 5 dpt (Two-way ANOVA, Tukey’s Multiple Comparisons Test). Data were averaged from at least 3 independent experiments. A minimum of 25 embryos were transfected in each experiment. Scale bar = 250 µm for (A) and 500 µm for (B).

### Embryo transfection is reproducible and consistent

To test the experimental reproducibility and consistency of the transfection protocol, we transfected 0.5, 1, and 2 µg of StayGold mRNA. We then measured the expression of StayGold in transfected embryos by measuring the fluorescence intensity in the whole embryo, and measuring gene expression with qPCR. To measure fluorescence intensity, we used the same acquisition settings in each condition and measured mean intensity in the whole embryo (Fig. 2A). We found that mRNA transfection yields consistent fluorescence intensity in each tested amount, and the fluorescence intensity values were consistent with the amount of mRNA transfected (Fig. 2B). To measure Stay-Gold gene expression, we pooled transfected fish for total RNA isolation and performed RT-qPCR. We confirmed that StayGold expression increased with increasing amounts of mRNA transfection and obtained consistent expression across experiments (Fig. 2C). Overall, transfection yields reproducible results without the need for laborious calibration.

**Fig. 2.**
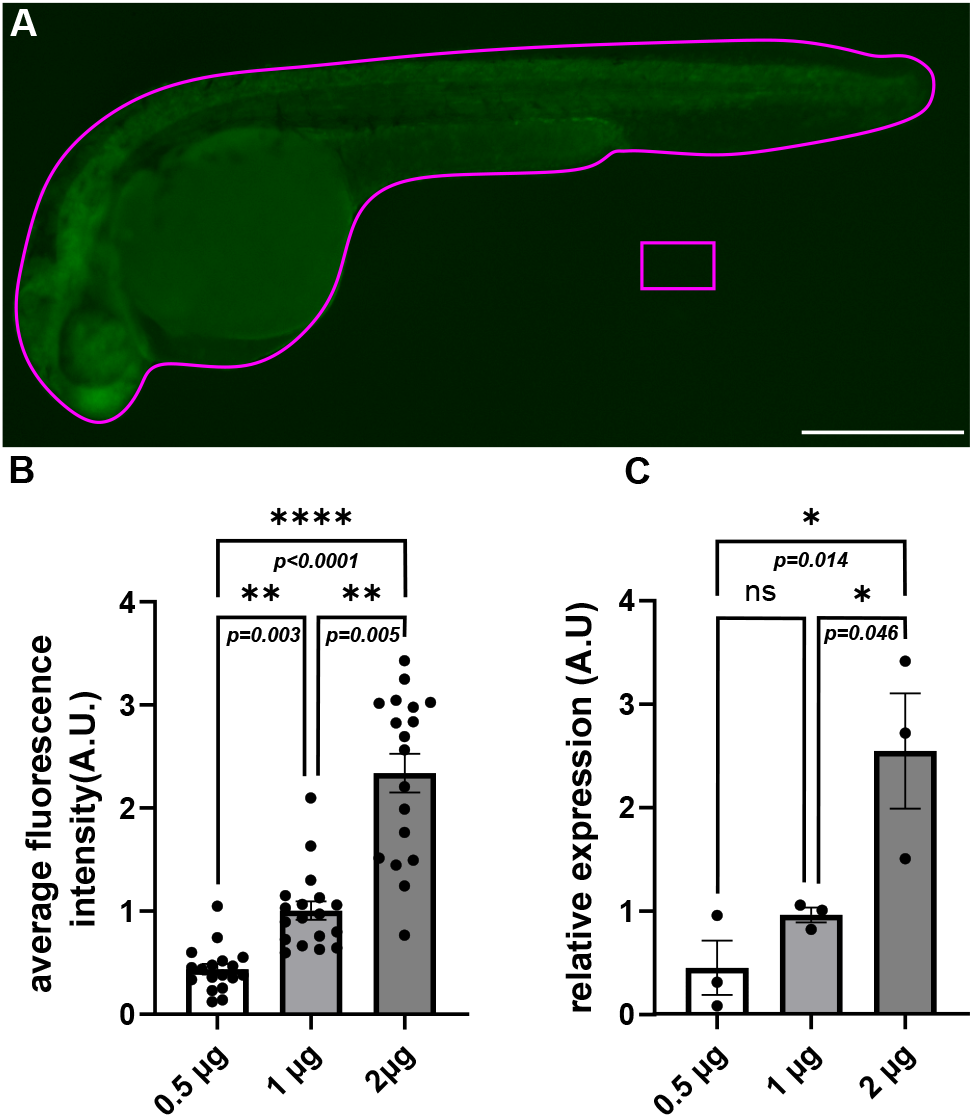
Transfection yields consistent and reproducible delivery of mRNA into zebrafish embryos. (A) An example image showing the ROI set for fluorescence intensity measurements. The average fluorescence intensity in each transfected fish was determined by subtracting the background (rectangular ROI) from the whole embryo ROI. (B) Bar graphs showing the average fluorescence intensity in embryos transfected with different amounts of mRNA. For each experiment, 6 embryos were measured per condition (Kruskal-Wallis Test, Dunn’s Multiple Comparisons). (C) Bar graphs showing the relative expression of StayGold mRNA assessed by RT-qPCR (One-way ANOVA, Tukey’s Multiple Comparisons Test). Data were acquired from three independent experiments. Each graph represents mean *±* SEM. Scale bar = 500 µm.

### Transfection allows for flexible incubation times and sparse labeling of embryos

To further study and refine the protocol, we tested different transfection conditions. First, we tested the minimum time required to keep embryos in transfection media to achieve successful transfection while not affecting survival. We found that leaving embryos in transfection media for 1 hour or 4 hours before exchanging media with 1X E3/methylene blue made no difference in the transfection efficiency (Fig. 3A) or the survival of embryos (Fig. 3B). Hence, transfection seems to occur in the first hour of embryos encountering Lipofectamine-mRNA complexes and embryos can be transferred to fresh plates with media exchange in a flexible time interval.

**Fig. 3.**
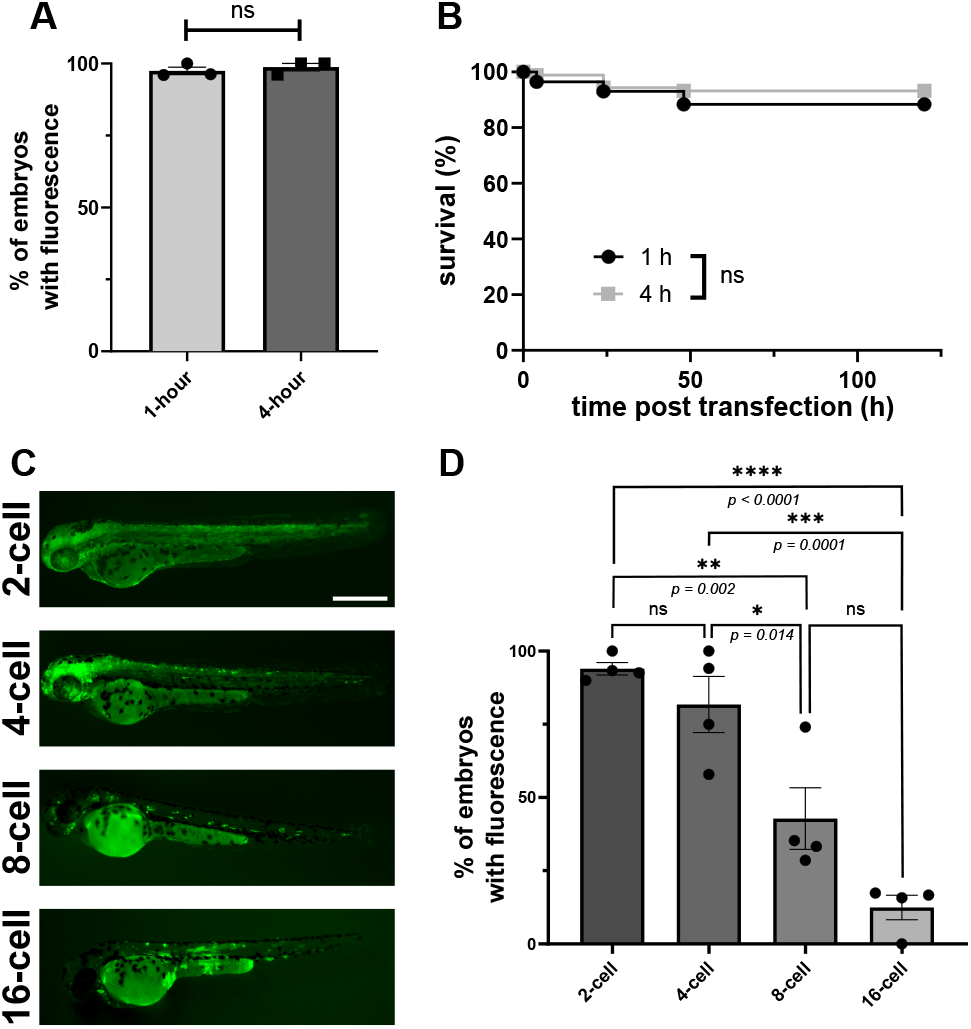
Our transfection protocol is flexible and allows for mosaic expression. (A) Bar graphs showing the percent of embryos with StayGold expression in 1- and 4-hour transfected embryos (Student’s t-test). (B) Survival of embryos over 5 days from transfection. 1-hour transfected embryo survival was not statistically different from 4-hour transfected embryo survival (Two-way ANOVA, Sidak’s Multiple Comparisons). (C) Images of 2 dpt embryos transfected with 2 µg StayGold mRNA at different cell stages. (D) Bar graphs showing the percent of embryos with StayGold expression when transfected at different developmental stages (One-way ANOVA, Tukey’s Multiple Comparisons Test). Data are averaged from at least three independent experiments. At least 20 embryos were transfected in each experiment. Data are shown as mean *±* SEM. Scale bar = 500 µm.

Next, we investigated whether transfection can occur in later stages of embryonic development. Occasionally, it is important to sparsely label cells for single-cell imaging and analysis. This is usually achieved by injecting after the first cell division to label a subset of cells. To test this, we transfected zebrafish embryos at two, four, eight, and sixteen-cell stages and compared the percent of embryos with fluorescence. We found that most of the embryos were still transfected at the two-cell stage, and transfection efficiency decreased gradually when transfected at later stages (Fig. 3C-D). While embryos transfected at the two-cell stage showed ubiquitous expression, later stages exhibited more sparsely labeled embryos (Fig. 3C). Overall, we found that lipofection is not limited to one-cell stage, and sparse labeling can be achieved by transfecting at later stages.

### Multiple Tol2 plasmids can be effectively delivered into zebrafish embryos in parallel with lipid-based transfection

Finally, we tested co-transfecting Tol2 plasmid with Tol2 mRNA (Kwan et al., 2007) for generating stable transgenic zebrafish lines with our transfection protocol. In these experiments, we also show that multiple constructs can be prepared and delivered in parallel, with minimal set-up overhead. We transfected three different Tol2 plasmids with the *eef1a1/1* (formerly known as *ef1a*) promoter (Negrutskii and El’skaya, 1998, Moon et al., 2013) driving expression of various nuclear fluorescent reporters, mNeon-Green, Cherry, and mCerulean (Fig. 4A). Tol2 plasmid-mRNA co-transfection did not affect the survival of embryos compared to control (Fig. 4B) and embryos developed normally. At 1 dpt, all three constructs exhibited fluorescence in transfected embryos (Fig. 4C-C”) and we detected persistent fluorescent signals at 7 dpt (Fig. 4D-D”). Across different experiments, we observed high transfection efficiencies (>80%) for all tested constructs (Fig. 4E-E”). The F0 larvae are being raised to screen for founders and, once ready, we will update this preprint to report on the germ line transmission efficiency. Overall, plasmids can be transfected efficiently and safely with transfection protocol, and parallel transfection offers a major advantage for simultaneously creating multiple transgenic zebrafish lines.

**Fig. 4.**
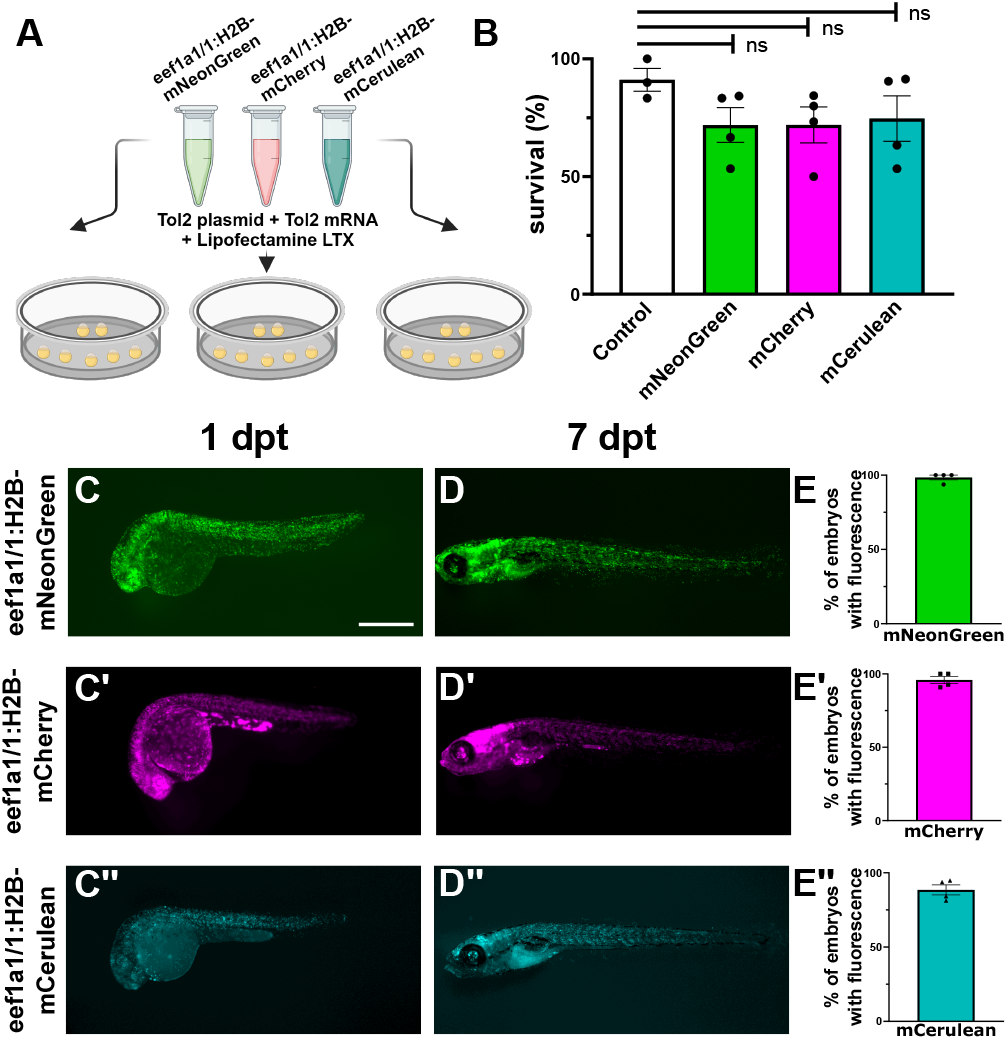
Parallel and efficient co-delivery of Tol2 plasmid and Tol2 mRNA into one-cell-stage zebrafish embryos with lipofection. (A) Schematic representation of the co-transfection protocol for Tol2 plasmid and Tol2 mRNA into zebrafish embryos. Multiple constructs can be transfected in parallel (B) Bar graphs showing the mean survival percentage of embryos co-transfected with different Tol2 plasmids at 5 dpt (*One-way ANOVA, Tukey’s Multiple Comparisons Test*. (C-C”) Fluorescence images of embryos transfected with *pT2-eef1a1/1:H2b-mNeonGreen, pT2-eef1a1/1:H2b-mCherry, pT2-eef1a1/1:H2b-mCerulean* at 1 dpt. (D-D”) Fluorescence of the transfected embryos at 7 dpt. (E-E”) Bar graphs showing the mean percentage of embryos with fluorescence. Graphs represent mean *±* SEM. Data averaged from at least three independent experiments. A minimum of 20 embryos were transfected in each experiment. Scale bars = 500 µm.

## Discussion

Here, we showed that lipofection can be efficiently used to deliver nucleic acids into zebrafish embryos. This protocol can be conducted without the need for special equipment and technical expertise, as opposed to microinjections. In this first version of our manuscript, we showed that the protocol is highly efficient in delivering mRNA and DNA, and that both types of nucleic acids can be co-transfected. As such, we expect this protocol to work for generating genetically modified fish lines using CRISPR (Jao et al., 2013, Medishetti et al., 2022, Mi and Andersson, 2023), or integrase-based methods (Lalonde et al., 2023). In the near future we plan to update our preprint to demonstrate some of these use cases. Our protocol is very flexible and open to modifications that can be applied for specific constructs without extensive optimization.

In addition, the protocol requires minimal training and it can be completed in less than an hour of work, massively reducing the amount of manual labor and time required for introducing nucleic acids into zebrafish embryos and for creating new transgenic lines.

The delivery of multiple different constructs using microinjection is a laborious and time-consuming process. Each construct needs to be loaded into a separate needle, which has to be then attached to a picopump. After that, each embryo is injected individually. Because of this, the number of separate constructs that can be injected by an individual during a session is limited to a handful. One major advantage of our protocol is that it can be highly parallelized. Multiple constructs can be prepared for independent transfection reactions with minimal overhead, and then delivered simultaneously to independent batches of embryos. Moreover, as most of the protocol requires only standard pipetting, it is very amenable to automation using liquid-handler robots. For these reasons, we believe this protocol offers great potential for use in high-throughput screens.

## ACKNOWLEDGEMENTS

We thank Dr. Akila Balasubramanian, Dr. Rachel Banks, Dr. Keir Balla, and Fitzgerald Small for their input in developing the protocol, and Dr. Sandra Schmid, Dr. Keir Balla, Dr. Akila Balasubramanian, and Dr. Emma Spikol for their comments on the manuscript. We also thank Dr. Valerie Tornini for teaching us embryo dechorionation. pCS2-Staygold plasmid was a gift from Dr. Keir Balla. The *eef1a1/1* promoter plasmid was a gift from Dr. Dan Wagner. The *pT2-eef1a1/1:H2b-mNeonGreen, pT2-eef1a1/1:H2b-mCherry*, and *pT2-eef1a1/1:H2b-mCerulean* plasmids were a gift from Dr. Xiang Zhao. Schematics were created with Biorender.com.

## Suplemental Information

### Zebrafish Embryo Transfection Protocol

#### Material List

| Reagent or Resource | Source | Identifier |
| --- | --- | --- |
| Pronase, Powder, Streptomyces griseus | Sigma-Aldrich | Cat.#10165921001 |
| Lipofectamine LTX Reagent with Plus<sup>TM</sup> Reagent | ThermoFisher | Cat.#15338100 |
| OptiMEM | ThermoFisher | Cat.#31985062 |
| Methylene Blue | Sigma-Aldrich | Cat.#03978 |
| Zebrafish AB strain | ZIRC | Cat.#ZL-1 RRID:ZIRC_ZL1 |
| Glass Petri Dishes-30mm | Amazon | Cat. #B09LGT1CGH |
| Glass Petri Dishes-60mm | Fisher Scientific | Cat.#08-748A, 316060BO |
| Leica Thunder Imager M205 FCA Microscope | Leica |  |
| Dumont #5 Forceps - Biologie/Dumostar | Fine Science Tools | Cat.#11295-10 |
| Glass Beakers | Fisher Scientific | Cat.#S63266 |
| Borosilicate Glass Pasteur Pipettes | Fisher Scientific | Cat.#NC0230004 |

#### Preparation of reagents

##### 1. Pronase

1.1 Prepare a 10 mg/mL stock solution by dissolving pronase powder in H_2_O.
1.2 Aliquot and store at -20 °C.

##### 2. 1X E3

2.1 Prepare a 60X stock solution by dissolving the ingredients in 2 L H_2_O.

2.2 Adjust pH to 7.2.
2.3 Dilute the 16.5 mL of 60X stock in 1 L distilled water to make 1X E3 medium.
2.4 Aliquot 500 mL of 1X E3 in a squirt bottle and add 50 µL of 1% methylene blue.

| 60x E3 Medium (2L) |  |
| --- | --- |
| NaCl | 34.8 g |
| KCl | 1.6 g |
| CaCl | 5.8 g |
| MgCl | 9.78 g |

#### Protocol: Step-by-step instructions

##### Day before transfection

###### 1. Fish set up and glassware preparation

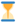 **Estimated time: 15 min - 2 h**

1.1 Incubate ∼ 50 mL of 1X E3 in an embryo incubator (28 °C) overnight.
1.2 Set up adult fish for breeding by separating male and female zebrafish in a breeding tank.
1.3 Autoclave glass dishes if necessary.
1.4 Flame polish the glass pipettes if necessary.

##### On the day of transfection

###### 1. Preparation of transfection reagents

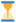 **Estimated time: 15 min**

1.1 Thaw pronase stock solution in a 37 °C water bath.
1.2 Prepare the pronase working solution by adding 0.8 mL of 10 mg/mL pronase stock solution in 15.2 mL pre-warmed 1X E3. Store in the incubator until ready for use.
1.3 Prepare 1 L and 250 mL beakers with 350mL and 150 mL 1x E3 for washes, respectively.
1.4 Prepare the nucleic acid-Lipofectamine mix as follows:

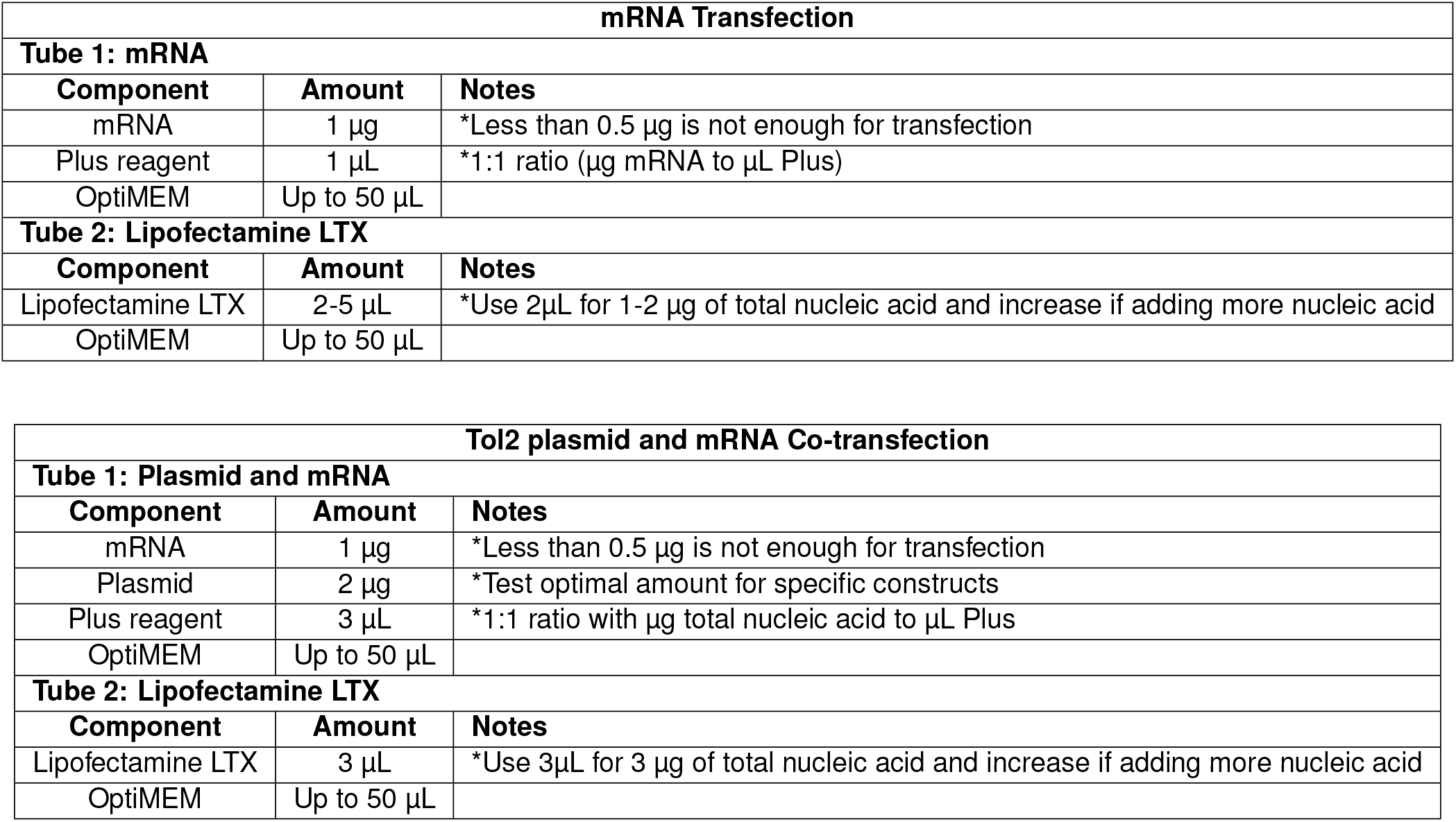
1.5 After adding Plus Reagent™ to the nucleic acid in Tube 1, gently flick the tube to mix and incubate for 5 min at RT.
1.6 Pipette Tube 1 into Tube 2, finger flick to mix.
1.7 Incubate at room temperature for at least 5 minutes. 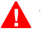 *The reagents can be incubated for up to one hour without affecting transfection efficiency*.
1.8 Add 400 µL of 1X E3 without methylene blue just before adding the mix to zebrafish embryos.

###### 2. Preparing zebrafish embryos for transfection

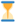 **Estimated time: 30 min**

2.1 Remove the divider between female and male zebrafish to allow breeding.
2.2 Let the females lay eggs and collect the fertilized embryos in 1X E3.
2.3 Transfer up to 100 embryos to a 60 mm glass dish. 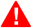 *Do not put more than 100 embryos per glass dish as this can lead to increased mortality*.
2.4 Remove all E3 and add ∼4-5 mL of pre-warmed pronase working solution.
2.5 Under a dissection scope, watch the embryos and use a hair-kife or pipette tip to squeeze chorions gently to check for wrinkling. Once 4-5 embryos show wrinkling in their chorions upon gentle squeezing, start washes. It usually takes 1-2 minutes for chorions to soften.
2.6 Allow embryos to settle in a 1L glass beaker filled with ∼ 350mL 1X E3.
2.7 Gently tilt the beaker to decant the 1X E3, allowing embryos to slowly migrate to the tilted side. Always leave enough media in the beaker to fully cover the embryos. 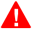 *Do not allow embryos to come in contact with air*.
2.8 Pour ∼150 mL fresh 1X E3 from the 250 mL beaker to wash the embryos. Repeat this wash step for 4 times. 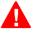 *At this stage, embryos should be out of their chorions. If needed, additional washes can be done to remove chorions from the remaining embryos*.
2.9 Gently collect the dechorionated embryos using a flame-polished glass pipette and place them in a 60 mm glass plate.
2.10 Transfer ∼30 dechorionated embryos in 30 mm glass dishes for transfection. 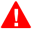 *Before moving the embryos into the glass dishes, add ample amounts of 1X E3. Make sure to have enough volume to completely cover the dechorionated eggs*.

###### 3. Transfection

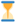 **Estimated time: 1 h - 4 h**

3.1 Remove as much 1X E3 as possible from the dechorionated embryos, first by pouring it and later using a pipette to remove the small remaining liquid. 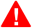 *Tilting the dish is helpful to remove media*.
3.2 Add 500 µL of the transfection mixture onto embryos.
3.3 Gently swirl the plate to mix.
3.4 Incubate for 1 to 4 hours in the embryo incubator at 28 °C. 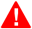 *Do not leave embryos in transfection media overnight*.
3.5 Change the media to 1X E3 with methylene blue, transfer embryos into 60 mm glass plates with flame-polished glass pipettes, and return them to the incubator.
3.6 The next day, fish can be moved into regular plastic petri dishes.

#### Troubleshooting

| Problem | Possible Cause | Suggested Solutions |
| --- | --- | --- |
| Embryos are not released from chorions | Insufficient pronase treatment | Make sure to prepare the pronase in pre-warmed E3 and keep in the incubator until use.<br>Increase the time of pronase treatment<br>Pipette up the embryos and slowly release them to break the chorions (mechanically) |
| High embryo mortality. | • Pronase carry-over<br>• Harsh pipetting<br>• A few dead embryo/debris contaminating the 1x E3 media<br>• Embryos left in transfection media for too long | • Decrease pronase treatment time<br>• Be more gentle while transferring dechorionated embryos, make sure to use glass and flame-polished pipettes<br>• Avoid placing dead embryos in the transfection dish<br>• Autoclave the glass dishes before transfection to avoid contamination from previous experiments<br>• Embryos need to be transferred to E3 with methylene blue after the transfection for survival |
| Underdeveloped embryos | • Embryo number per transfection volume is too high | • Decrease the number of embryos per dish<br>• Increase the volume of transfection media<br>• Transfer embryos in a bigger dish after transfection |
| No transfection | • Nucleic acid cannot combine with the Lipofectamine LTX | • Use 1X E3 without methylene blue in transfection media.<br>• Make sure mRNA/DNA is intact and not degraded by checking on agarose gel or tapestation, if available.<br>• Make sure OptiMEM is not contaminated. Use fresh aliquots in each experiment and keep OptiMEM at 4 °C.<br>• Increase the incubation time of nucleic acid-LTX to 30 min - 1 h. |
| Decreased transfection efficiency | • Liposomes are diluted out in larger volumes<br>• Nucleic acids are degraded<br>• Liposome-nucleic acid formation is not complete | • Adjust the volume of transfection and/or scale up the nucleic acid-Lipofectamine amount<br>• Work in a clean area. Use RNase Out if necessary<br>• Increase the time of incubation of Lipofectamine LTX and nucleic acid |
| Embryos dissociating after transfection | • E3 was not added in the transfection media after liposomes were mixed with the nucleic acid | • Make sure to dilute LTX-OptiMEM in E3 while transfecting. Do not leave embryos in only LTX-Optimem media |

